# Carbon dots deposition in adult bones reveal areas of growth, injury and regeneration

**DOI:** 10.1101/2020.10.13.338426

**Authors:** Rachel DuMez, Esmail H. Miyanji, Lesly Corado-Santiago, Bryle Barrameda, Yiqun Zhou, Sajini D. Hettiarachchi, Roger M. Leblanc, Isaac Skromne

## Abstract

C-dots synthesized from carbon nanopowder (oxidation, hydrothermal) are particularly attractive theragnostic agents for bone-related injuries and disease due to their bright fluorescence and high binding affinity and specificity for bones, as demonstrated in a larval animal model. Larval bone development, however, is significantly different from the bone growth, repair and regeneration processes occurring in adults. Using adult zebrafish, we investigated C-dots’ interactions with adult skeletal structures. Upon injection, C-dots were observed at the surface of bones, at sites of appositional growth. In regenerating bones, C-dots were observed at the core and on the surface of the bones depending on the age of the tissue. C-dot’s deposition occurred within 30 min of delivery and it was highly selective. Importantly, their deposition did not interfere with bone regeneration or the animal’s health. Together, these properties establish C-dots as novel tools for the diagnostic and treatment of adult bone-related injuries and diseases.

---

Carbon-based nanoparticles or carbon dots (C-dots) have emerged as novel therapeutic and diagnostic biomaterials due to their unique, tunable physicochemical properties^1, 2^. Common to all C-dot species is their small size (<10nm), high carbon content, high photostability and bright fluorescence^2, 3^. Several factors contribute to these properties, most significantly, their core configuration and surface functional groups: core and surface molecular states and quantum confinement effects directly influence C-dot’s maximum excitation and emission wavelengths, excitation-dependent photoluminescence and fluorescent quantum yield^3–5^. C-dots’ core structure and surface functionality groups can be modified during their synthesis, making their photoluminescent properties highly tunable for a broad range of bioimaging applications^1, 2^. In addition, the ability to modify surface functional groups after synthesis can further expand C-dot’s utility beyond imaging, for example, by serving as pharmaceutical nanocarriers^6, 7^. The surface of C-dots can be passivated (e.g., Polyethylene glycol;^8, 9^) and conjugated covalently or noncovalently with diverse therapeutic drugs^10–12^. Among various surface modification strategies, formation of covalent bonds can have unexpected consequences for both the C-dot vehicle and the drug cargo, including changes in solubility, therapeutic efficacy, and systemic clearance of the drug^13–15^. Therefore, to fully exploit C-dot’s theragnostic potential, it is imperative to characterize unmodified C-dots.

Diseases impose heavy societal burdens that can be mitigated with improved diagnostic techniques and expanded treatment options. Bone diseases such as osteoporosis and low bone mass combined affect an estimated 54 million individuals annually in the U.S. alone^16^. Significantly, early diagnosis is key for the most successful treatment prognosis. Currently, treatments predominantly rely on preventing further bone erosion and not in restoring bone mass, as drugs that promote bone growth can lead to cell proliferation in other tissues and increase a patient’s cancer risk^17^. Diagnosis relies on x-ray imaging methods, MRI or CT scans, with novel fluorescent-based technologies of higher sensitivity currently under development^4, 18–20^. The development of more sensitive diagnostic tools for early bone-loss detection and novel treatment methods for stimulating bone growth without affecting other tissues will significantly ameliorate the societal and personal cost of bone diseases.

We recently described that C-dots synthesized from carbon nanopowder have a high affinity and specificity for developing zebrafish bones^21, 22^. Zebrafish provides a robust, *in vivo* model to test C-dots’ interaction with biological tissues, as their transparency allows direct observation of C-dot’s photoluminescence. More importantly, bone development in zebrafish is remarkably similar to that of mammals^23–25^. In all vertebrates, bones develop either through direct aggregation of bone-forming cells (intramembranous ossification; e.g., cranial bones) or through the deposition of mineral matrix on a collagen scaffold (endochondral ossification; e.g., long bones)^26^. Once formed, adult bones undergo homeostatic turnover and can continue to increase in diameter through the process of appositional growth, whereby new bony tissue is added to the bone’s surface^26–28^. When carbon nanopowder derived C-dots were injected into 5 day old zebrafish larvae, their deposition was observed in opercular (intramembranous) and in vertebrae (endochondral) bones, but not in non-skeletal tissues^21, 22^. Their deposition on bones was dependent on bone mineralization, as manipulations promoting or interfering with bone mineralization increased or decreased C-dot deposition, respectively^21^. Importantly, only C-dots synthesized from carbon nanopowder (oxidation; hydrothermal) and rich in surface carboxyl groups bind to bones, whereas C-dots synthesized using other approaches (e.g., citric acid and EDA; solvothermal) and rich in surface amine groups did not display this property^22^. To date, all analysis of C-dot deposition on bones was done in larvae. Whether C-dots also bind to adult bones undergoing homeostatic turnover and appositional growth remains unknown. Given that osteoporosis, other bone related diseases and trauma primarily affect aging individuals, it is important to characterize the activity of C-dots in an adult animal model.

To gain insight into the adult theragnostic potential of carbon nanopowder-derived C-dots, we analyzed their toxicity, bone binding dynamics and photoluminescence in transparent adult zebrafish (Casper strain;^29^). We analyzed C-dot’s bone deposition using their intrinsic photoluminescence (excitation peaks between 360-540 nm and emission between 500-600 nm^15^) and a standard 488/525 nm excitation/emission Fluorescein filter set. Three different delivery methods were used to determine the ability of C-dots to bind adult bones, one cutaneous and two through injection (Fig. 1). To determine if C-dots can be adsorbed cutaneously, we applied a concentrated solution of C-dots to the caudal fin or the flank of the fish. Neither application resulted in labeling of local bones (Fig. 1A, B, and data not shown). Similarly, topical application of C-dots to a healed caudal fin wound undergoing regeneration did not label local bones (Fig. 1C, D). In contrast, C-dots applied to a wound with exposed bones labeled the exposed tips of the bones (Fig. 1E, F). Notably, none of the bone tissues that regenerated following C-dots exposure was labeled, indicating that following deposition, C-dot binding became fixed. We next employed two injection delivery methods, intraperitoneal and intravascular (retro-orbital). Delivery of C-dots (20 *μ*g/g fish) through either method effectively labeled intact adult bones (Fig. 1G-O). This labeling was most apparent in the fish fins because of the tissue’s reduced thickness and isolated position (Fig 1G-O, R, S). These results indicate that both injection methods are effective for delivering C-dots into adult fish and, consistent with our previous reports in larvae^15^, that C-dots can bind with high specificity to skeletal elements regardless of bone age.

**Figure 1.**
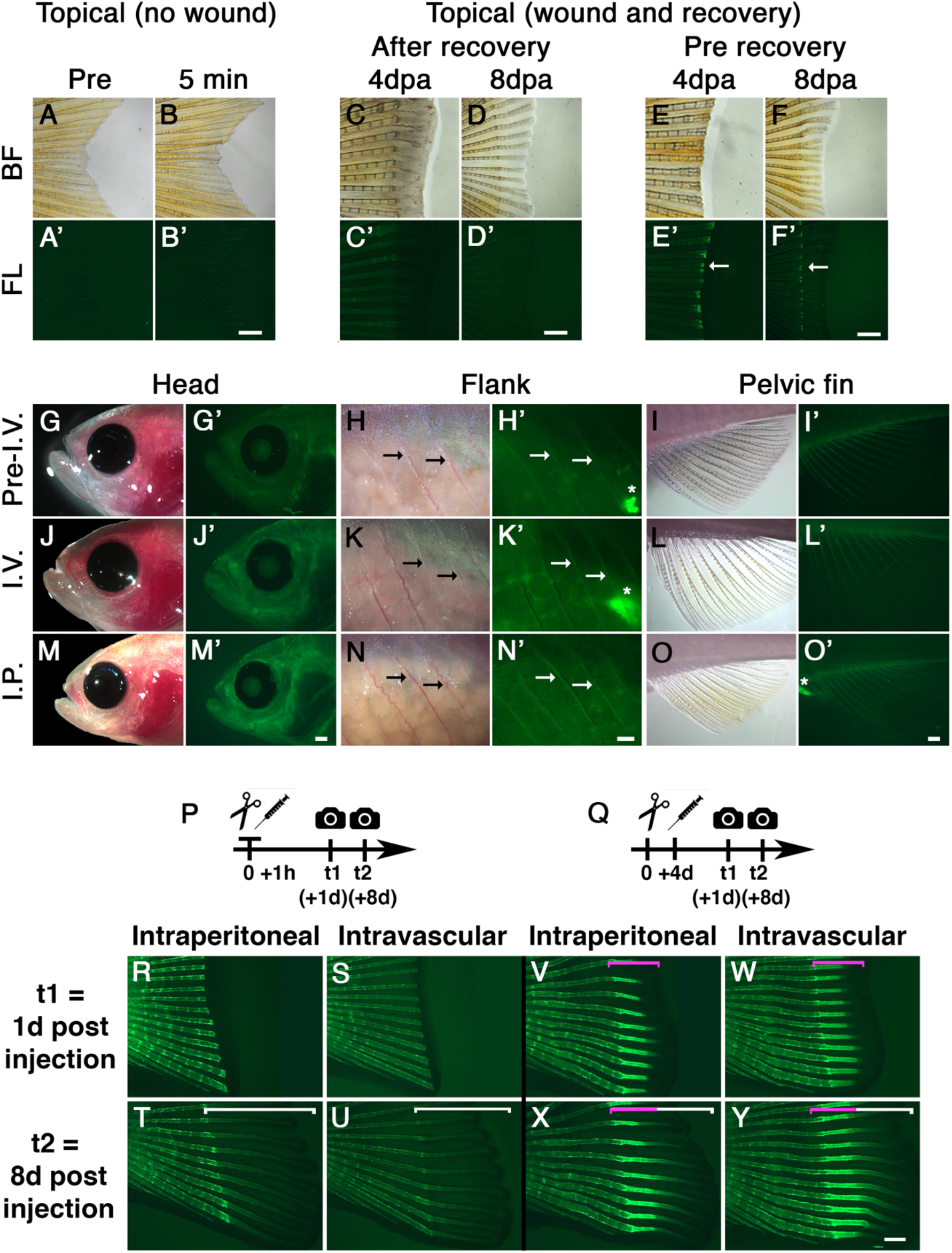
C-dots binding to adult ossified tissue. **(A, B)** Topical exposure of C-dots to skin (2 min; 2*μ*g/ml) do not label local bones fluorescently. Images in B were taken 5 min after C-dots exposure. **(C, D)** Topical exposure of C-dots to a healed wound (caudal fin amputation, 4 and 8 day healing and regeneration), does not label local bones. **(E, F)** Topical exposure of caudal fins to C-dots immediately after amputation resulted in bone labeling at the wound site (green fluorescence; white arrow). Bones that regenerated after healing were not labeled with C-dots. **(G-I)** Images of head, flank and pelvic fin of fish prior to intravascular injection of C-dots under reflected bright field (BF) and fluorescent (FL) illumination. Thick tissues naturally display a low level of background fluorescence. **(J-L)** Intravascular injection (I.V.) of C-dots resulted in fluorescent labeling of ossified tissue. **(M-O)** Intraperitoneal injection (I.P.) of C-dots also resulted in fluorescent labeling of bones. **(P, Q)** Schematics showing timeline of experiments. C-dots were injected before (P) or after (Q) the onset of regeneration and their deposition on bones was analyzed on day 1 and 8 post injection. **(R-U)** Before the onset of regeneration, injected C-dots deposit only in non-regenerating bones. Brackets indicate areas of bone regeneration. **(V-Y)** After bone regeneration has been initiated, injected C-dots deposited in non-regenerating and regenerating bones. Brackets indicate areas of bone regeneration, with magenta domain indicating areas of regeneration between days 4 and 5 after injection. n=6 fish per experimental condition, in two independent experiments. Time 1 and 2 images are of the same fish. Fish are positioned anterior to the left and dorsal to the top. In flank images, arrows indicate ribs and asterisks nonspecific gut autofluorescence. Day post amputation is indicated as dpa. Scale bars are 500 *μ*m.

To further explore the binding of C-dots to adult bones, we administered C-dots to adult zebrafish that had undergone amputation of the caudal fin and were at different stages of regeneration. This approach allowed us to examine the binding of C-dots to fin ray bones undergoing normal homeostatic turnover as well as regenerative growth. When C-dots were delivered prior to the initiation of fin bone regeneration at 1 day post amputation (dpa), regenerated bones examined at 8 dpa were not labeled (Fig. 1P, R-U). However, when C-dots were delivered during active fin bone formation (e.g., 4 dpa), regenerated bones were strongly photoluminescent (Fig. 1Q, V-Y). The observation that C-dots delivery at 4 but not 1 dpa labeled regenerating bones at 8 dpa suggests that C-dots were quickly cleared from circulation. Attempts to determine if C-dots were metabolized or excreted were unsuccessful, as C-dot concentrations were below the detection limit of our assays (ELISA and TEM; data not shown). Therefore, at this moment, we cannot rule out the possibility that C-dots were removed from the circulatory system through mechanisms other than bone deposition. Nonetheless, once in bones, C-dots label could be detected for several weeks (Fig. 1; and data not shown), suggesting that binding to bones is permanent. In terms of delivery method, intravascular injections gave more consistent results than intraperitoneal injections, as C-dots distribution through blood vessels was immediate and avoided their sequestration in cavities such as the peritoneum (data not shown). Together, these results suggest that intravascular injection is an effective C-dots delivery method and that C-dots bind to adult bones for long time periods.

The photoluminescence intensity of C-dots bound to regenerating bones was qualitatively more intense and homogeneous than the binding to non-regenerating bones, providing an assay to study C-dots binding to ossifying tissue. To begin testing the magnitude of C-dot binding to regenerating bone as a function of C-dots dosage, we repeated the amputation experiment followed by the intravascular administration of increasing amounts of C-dots. The concentration of C-dots delivered to each fish was normalized to the weight of the fish (*μ*g/g). The lowest amount at which we were able to detect C-dots binding to regenerating bones was 5 *μ*g/ml (Fig. 2A). The photoluminescence intensity increased until 40 *μ*g/g, the highest dose tested (Fig. 2A). Higher concentrations were not tested because concentrations above 40 *μ*g/g clogged the microneedles used for injections and the use of larger needles damaged the fish’s vasculature at the site of injection. To quantify C-dots’ binding, we measured the photoluminescence intensity of a 100 × 100 *μ*m square area located 100 *μ*m away from the amputation site in regenerating fin rays 4, 6, 8 and 10 (ventral to dorsal position). These values were averaged and normalized to the intrinsic background of corresponding fish as well as to control fish (0 *μ*g/g). The normalized averages were then plotted against C-dots dosages to determine the magnitude of binding. Based on three independent experiments, there is a linear relationship between C-dots dosage and photoluminescence intensity (Fig. 2B; linear regression, R^2^>0.96; Standard t-test, df=3, p<0.002). Trendlines did not plateau indicating that tissue saturation was not achieved. Together, these results suggest that concentrations of C-dots ranging from 5-40 *μ*g/g can label regenerating bone without compromising the fish’s health or the process of bone regeneration. Due to concern of potential tissue damage during injection of a high volume of C-dots, in subsequent experiments we used 20 *μ*g/g of C-dots.

**Figure 2.**
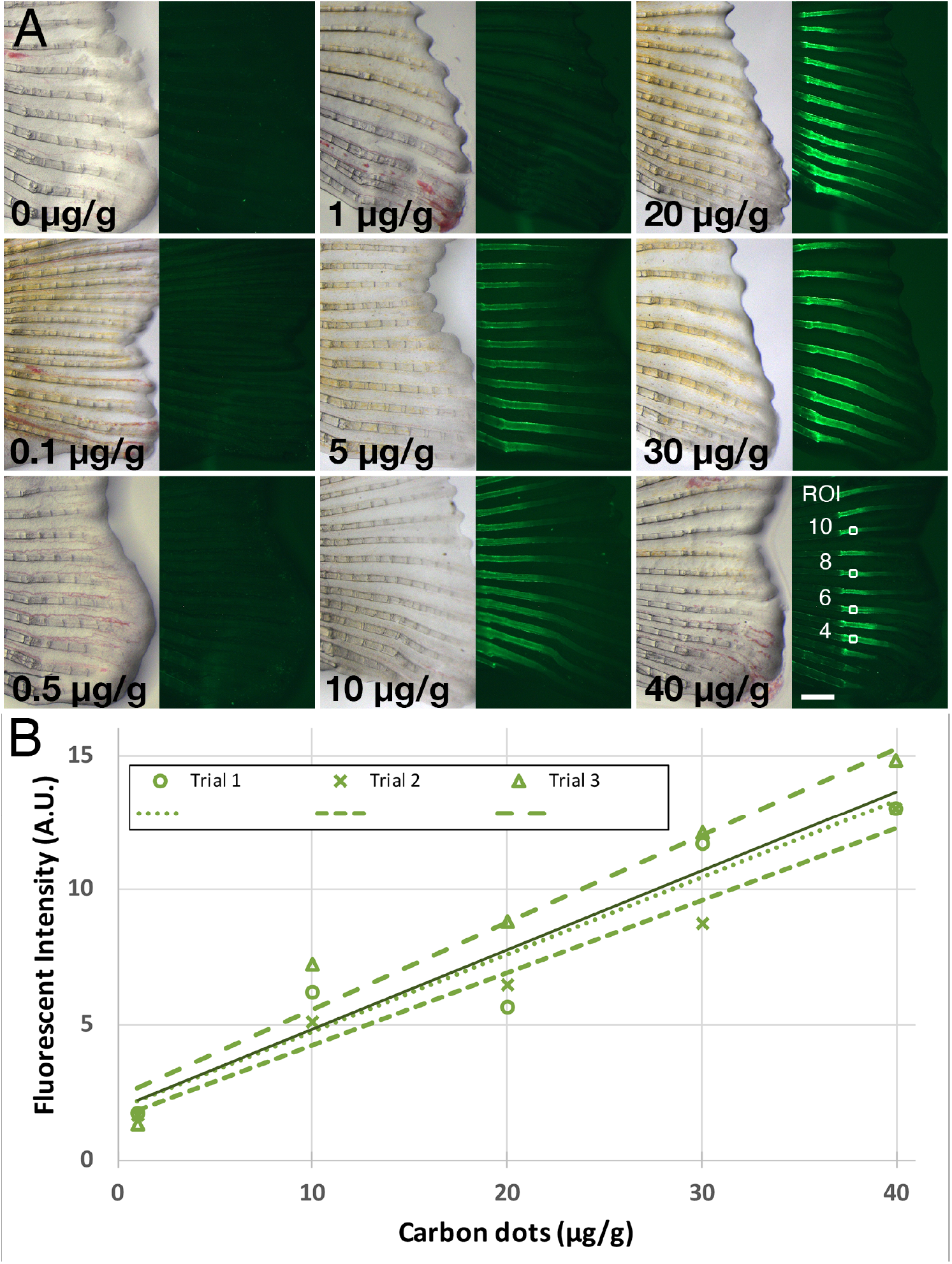
C-dots bind to regenerating bones in a concentration-dependent manner. **(A)** C-dots deposition on regenerated bones correlates with concentration of C-dots injected. Timeline of experiment as in Fig. 1Q, except that images were collected 4 days post injection. C-dot’s concentration was normalized to fish weight (*μ*g/g; 0 is saline-injected control). Representative regions of interest (ROI) used for fluorescence intensity quantification are indicated with squares for the 40 *μ*g/g injected fish fin. Scale bar is 500 *μ*m. **(B)** Mean fluorescent intensity of bone-bound C-dots at ROIs as a function of C-dots dosage. Values were normalized to background and saline-injected controls and reported in arbitrary units (A.U.). Dashed lines represent trendlines for three independent experiments (R^2^>0.91; df=3, p<0.006). Solid line represents the mean average trendline from the three experiments (R^2^>0.96; df=3, p<0.002).

In the previous experiment, C-dots were injected at a fixed time during fin regeneration. To determine whether C-dots differentially deposition on bones is dependent on the state of regeneration, we repeated the experiment varying the time of injection after amputation. Time of C-dot injection varied from 2 to 7 dpa, and all the regenerated fins were imaged at 8 dpa (Fig. 3A). We then quantified the photoluminescence intensity profile of the C-dots deposited in the fourth most ventral regenerated fin ray (Fig. 3B). Compared to controls, injection at 2 dpa resulted in the poor bone labeling least amount of C-dots labeling, and only at the site of amputation. Injection at subsequent days increased the area of labeling along regenerating bones; the whole regenerated ray was labeled at 7 dpa (Fig. 3A). Signal quantification of the labeled bones revealed an inverse correlation between the amount of tissue labeled and the photoluminescence of the labeling: the more area labeled, the lower the intensity of the photoluminescent signal (Fig. 3B). These observations suggest that C-dots deposition after fin amputation occurs at all stages of bone regeneration, resulting in an apparent homogeneous distribution of fluorescent labeling throughout the bones. The clearance of C-dots from circulation in the regeneration assay was swift, with C-dot depositing on regenerated bone at the time of delivery, and not on bone that regenerated after that time. For example, injection at 3 dpa only labeled the bone that had regenerated at that point, and not bone that regenerated between 4-7 dpa (Fig. 3 at 3dpa). Together these results suggest that C-dots bind to available regenerated fin ray bones at the time of injection.

**Figure 3.**
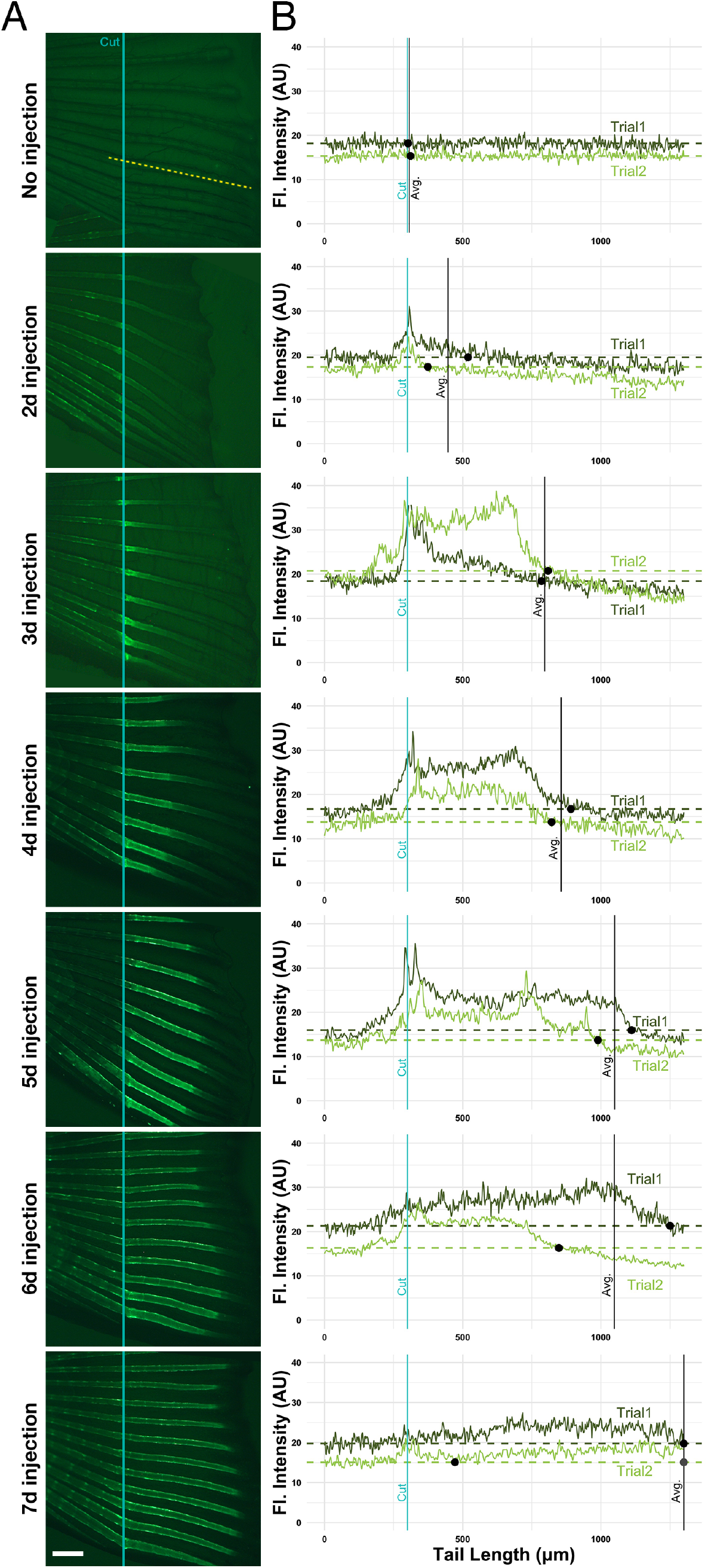
C-dots distribute evenly throughout available regenerating bones. **(A)** C-dots deposit on available regenerating bone at time of injection. Injection days after amputation are indicated on the left. Specimens were imaged 8 days after amputation, anterior to the left. Site of the cut is indicated with a blue line, and the site of the profile intensity analysis with a yellow dashed line. Experiments were done in duplicate, with three fish per trial. Scale bar is 500 *μ*m. **(B)** Fluorescent intensity quantification profile across the length of the fourth ventral fin ray (bone), in two representative fish from two independent experiments. Fluorescent signal in arbitrary units (A. U.) was normalized to saline-injected controls. Blue line indicates the site of amputation, black dots indicate the position where fluorescence in regenerating bone reaches average background fluorescence levels (non-regenerated portion of the bone; dashed line), and black line indicates the position where average fluorescence levels in both specimens reach background levels (except in 7-day fish, were average for trial 2 fish reaches background levels twice; gray dot).

To investigate the dynamics of C-dot deposition at the tissue level, we analyzed the distribution of C-dots’ photoluminescence in transverse sections of regenerating caudal fins. Adult zebrafish were injected with C-dots 1, 4 or 8 dpa and, after 1-hour of labeling, the fin was harvested, cryosectioned, and imaged using confocal microscopy. We did not observe any histological differences in the regenerated bones between control and C-dot injected fish (Fig. 4 and data not shown), supporting the gross morphological observations that C-dots did not interfere with bone regeneration processes. At all stages of regeneration, bones proximal to the amputation site were thicker, more mineralized, and exhibited more intense C-dot signal than bone located at the fin’s distal tip (Fig 4B). To test if bone age is a determinant of C-dot deposition, we analyzed cross sections of bones at different stages of regeneration at comparable proximal-distal positions. Comparison of representative sections adjacent to the amputation site revealed weak labeling of newly regenerated bones at 1 dpa, strong and even labeling at 4 dpa, and strong labeling of bone surfaces at 8 dpa (Fig. 4). Given that the diameter of bones increases by the repeated addition of ossified tissue at the surface (appositional growth^27^), our observations suggest that C-dot deposition occurs in areas of bone growth.

**Figure 4.**
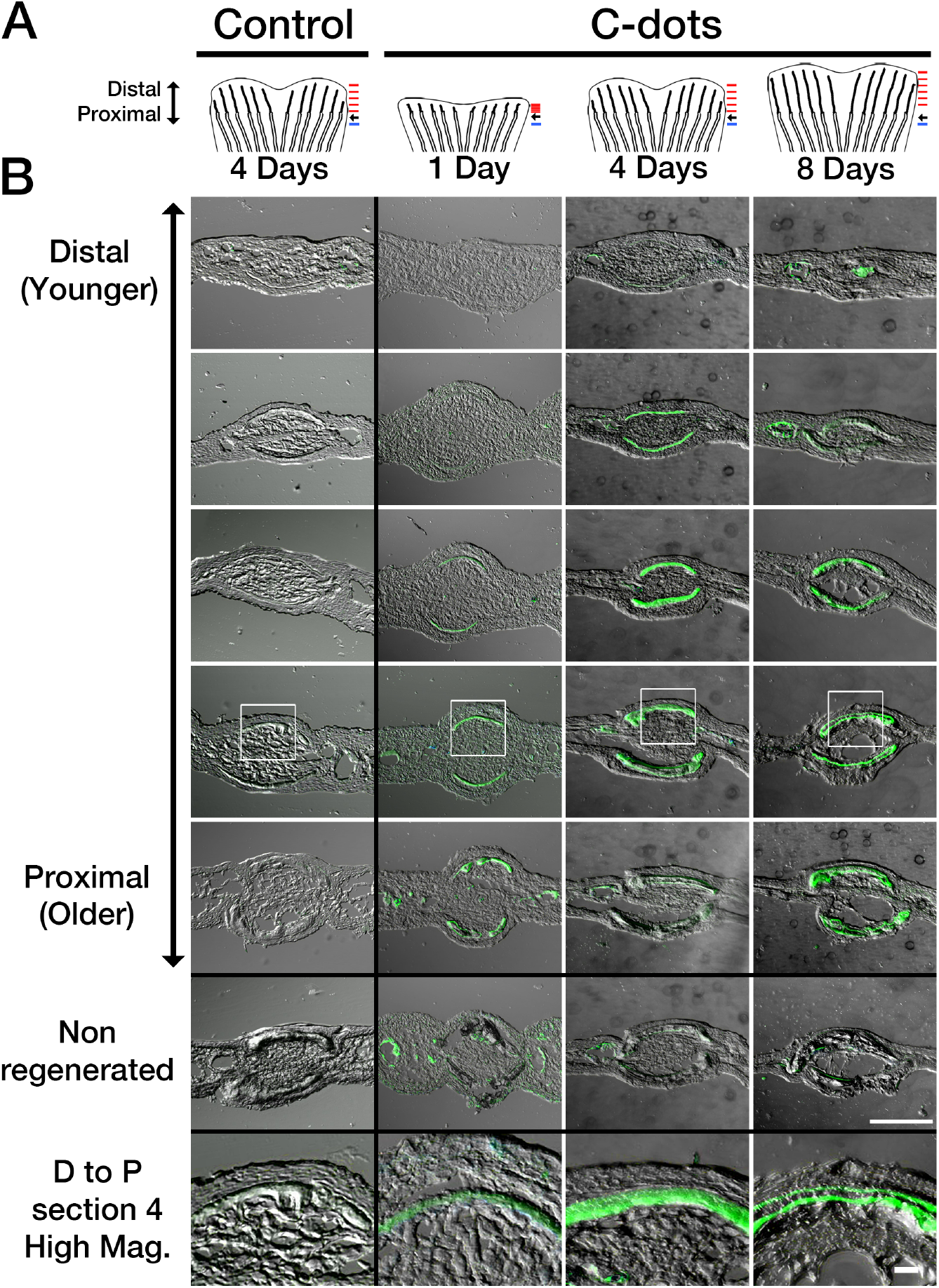
Differential spatiotemporal deposition of C-dots in regenerating bones. **(A)** Diagram of regenerating fins used in analysis. Amputated fish were injected with saline (control) or C-dots at the indicated times. One hour after injection, fins were re-amputated and processed for cryosectioning. Black arrow indicates amputation site. Red and blue lines indicate the approximate site of sections in regenerating and non-regenerating bones, respectively. **(B)** Sagittal sections of regenerated and non-regenerated caudal fins ordered in distal (younger) and proximal (older) direction (position corresponding to red and blue lines in A, respectively). Areas of C-dot deposition are green. Areas in white box are shown magnified in the bottom panels. Scale bar is 100 *μ*m for low and 20 *μ*m for high magnification images.

To directly test C-dots deposition in areas of tissue growth, we examined the distribution of C-dots relative to that of the dye Alizarin Red Complexone (ARC). Intravital staining of bones by Alizarin red dyes is a well-established histological method used in mice to distinguish areas active bone growth, as the dye is preferentially taken up by these areas compared to regions where the bone has stopped growing^30^. Thus, co-localization of C-dots and ALC signals would indicate that C-dots preferentially bind to areas of active bone growth. To validate the use of ALC in zebrafish, we exposed fish undergoing caudal fin regeneration to ALC at 8 dpa. In these fish, regenerated fin bones were strongly stained with ALC (Fig. 5A). In cross sections, ALC’s stain distribution was homogeneous in bones closer to the fin tips (Fig. 5D, D’), and superficial in bones closer to the amputation (Fig. 5E, E’). These findings are similar to those observed during normal growth of bones in mammals^30^, supporting the use of ALC as a viable method for identifying areas of bone growth in zebrafish.

**Figure 5.**
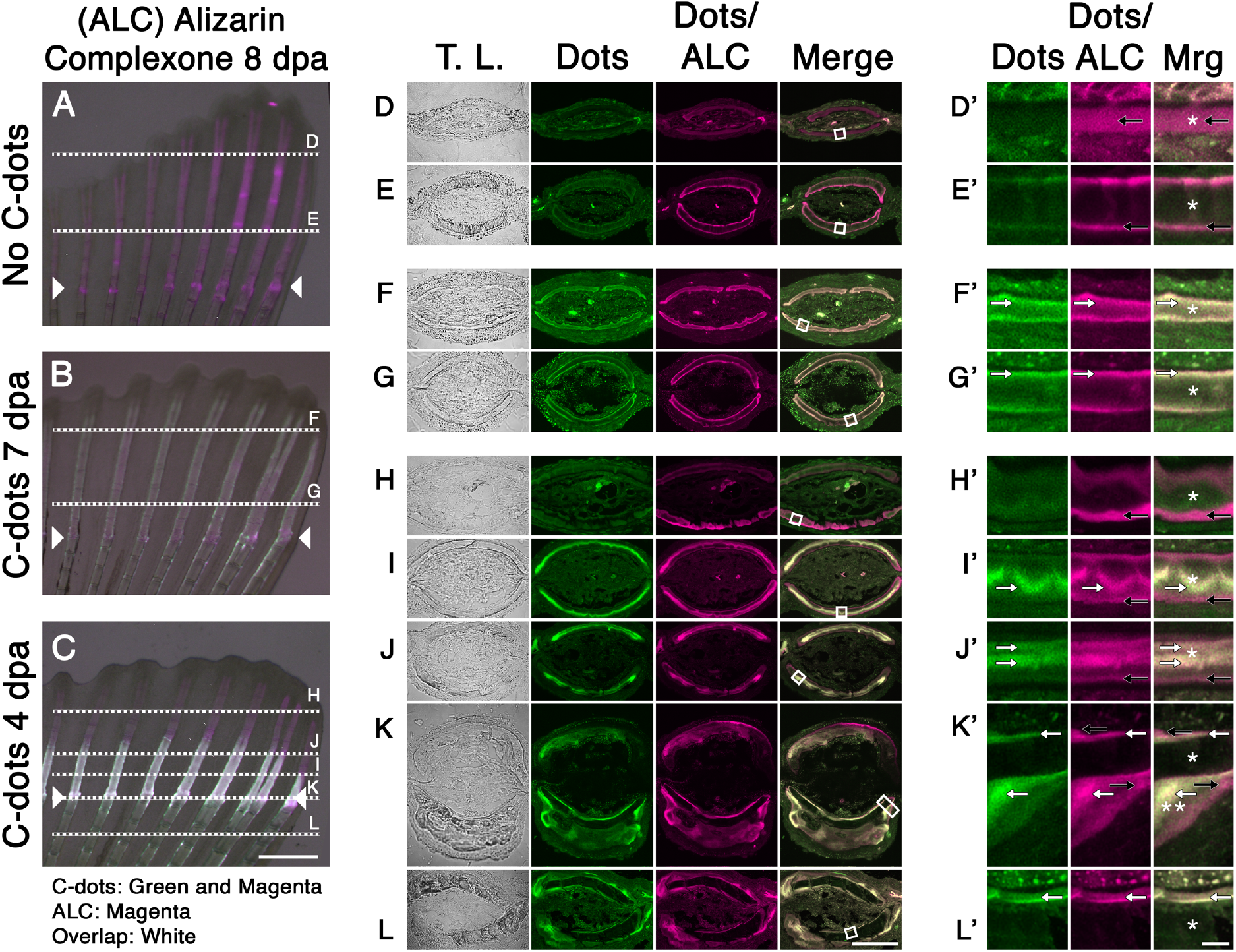
C-dots deposit at sites of active bone mineralization. **(A-C)** Regenerated caudal fin of uninjected fish (A), injected with C-dots at 7 dpa (B), or injected at 4 dpa (C). All fish were stained with Alizarin Complexone (ALC) at 8 dpa and imaged. Composite images were generated by superimposing photographs obtained using transmitted light, a 488/525 nm filter set to capture C-dot’s fluorescence (pseudo colored green), and a 596/615 nm filter set to capture C-dots/ALC’s fluorescence (pseudo colored magenta). ALC-positive areas (indicative of ossification) that lack C-dots appear magenta, and areas of ossification with C-dot deposition appear white. Posterior end of fin towards top of images. Site of amputation is indicated with arrowheads and the plane of sections D-L are indicated with dotted lines. **(D-L)** Transverse sections of fins, imaged and pseudo colored as in A-C. **(D’-L’)** Magnified bone regions from D-E (white squares). Areas of C-dot deposition are indicated with white arrows. Ossified regions devoid of C-dots are indicated with black arrows. Asterisks indicate the bone matrix. Double asterisk in K’ indicates areas of matrix buildup at the site of amputation. Scale bars are 500 *μ*m for A-C, 50 *μ*m for DL and 5 *μ*m for D’-L’.

Next, we investigated the relationship between C-dot deposition and areas of ALC staining. To identify areas of ALC staining, it was necessary to use an indirect analysis method, as C-dot’s 500-600 nm emission range overlaps with the emission peak of ALC (580 nm). Areas of growth devoid of C-dots were identified by the presence of signal at 580 nm (red filter; C-dots and ALC) but not at 525 nm (green filter; C-dots only). To begin exploring C-dot deposition in areas of active bone growth, we injected fish with C-dots at 7 dpa and stained them with ALC at 8 dpa (Fig. 5B). Because bone layer formation takes more than twenty-four hours, we expected to see C-dots and ALC signal co-localization. In all sections examined, C-dots and ALC signals overlapped (Fig. 5F, F’, G, G’). At the distal tips of the fins, where new bone was being generated, both stains were observed throughout the bone (Fig. 5F, F’). In areas closer to the amputation site, both signals were confined to the surface of the bones (Fig. 5G, G’). This extensive signal overlap precluded the unambiguous identification of ALC positive areas of bone growth devoid of C-dots. To overcome this challenge, we spatially separated temporal areas of growth by injecting fish with C-dots at 4 dpa and staining them with ALC at 8 dpa. In this experiment, we predicted that areas of early bone growth would be positive for C-dots but not ALC, whereas areas of late bone growth would be positive for ALC but not C-dots. Analysis of whole fins revealed ALC staining throughout the length of the bones, while C-dots labeling was restricted to the proximal portion of bones (Fig. 5C). The distal sections of fin bones that hadn’t formed prior to the injection of C-dots (4 dpa) were negative for C-dots but positive for ALC (Fig. 5H, H’). Similarly, bone tissues close or proximal to the site of amputation had C-dots deposits and ALC staining at the bone surface (Fig. 5K, K’, L, L’). Strikingly, bones in the process of regeneration at the time of the injection showed a dual staining pattern: the core portion of the bone showed C-dot deposits while the surface of the bone was ALC positive (Fig. 5I, I’, J, J’). Taken together, these findings suggest that C-dots bind to areas of bone growth.

In summary, here we demonstrate that in zebrafish, C-dots derived from oxidized carbon nanopowder can bind to adult bones in areas of tissue growth. We observed C-dots’ deposition at the surface of bones undergoing homeostatic turnover, at sites of acute trauma (fracture), and in areas of regeneration (Fig. 1), in a manner proportional to the quantity of C-dots injected (Fig. 2, 3). All these sites are regions where the bone is actively growing, as determined by the colocalization of C-dots and Alizarin red, a commonly used dye that labels regions of bone growth (Fig. 5;^30^). These results build on our previous findings in developing zebrafish larvae bones^21, 22^. Importantly, after intravenous delivery, C-dots are cleared from the circulation within 30 min and are found evenly distributed across all bones in the body (Fig. 1). The binding of C-dots to bones appears to be permanent; after the original injection, C-dots can be detected in bones for several weeks (Fig. 1) and up to several months (data not shown). These observations, together with the fact that C-dot deposition does not appear to interfere with bone repair and regeneration strongly suggest that C-dots are inert. These findings have important implications for future applications of C-dots as a drug delivery vehicle for the treatment of skeletal diseases and traumatic bone injuries.

## ASSOCIATED CONTENT

### Supporting Information

Materials and Methods (Microsoft Word)

R-script C-dots fluorescent quantification (PDF)

## Author Contributions

The manuscript was written through contributions of all authors. All authors have given approval to the final version of the manuscript.

## ACKNOWLEDGMENT

We want to thank the members of the Leblanc and Skromne lab for encouragement and support. We also want to thank members of the Lambert lab for training and the use of the cryostat; Christie Lacy for help with the confocal microscope; Dr. Omar Quintero for reagents and advice on fluorescent image analysis; and Dr. Kristine Nolin for the use of her chemistry laboratory.

## ABBREVIATIONS

C-dots: carbon nanodots
dpa: days post amputation
dpi: days post injection
ARC: Alizarin Red Complexone
ELISA: Enzyme-linked immunosorbent assay
TEM: transmitted electron microscopy

## SUPPLEMENTAL INFORMATION

### Materials and Methods

#### Carbon Dots Synthesis

Carbon nanodots (C-dots) were synthesized from carbon nanopowder and purified using our previously reported procedure^1, 2^. To obtain C-dots in powder form, the C-dot solution was lyophilized in a rotovap evaporator at 60°C. The morphology and photoluminescent properties of as-prepared C-dots were confirmed using transmitted electron and epi-fluorescence microscopes, as previously described^1, 2^.

#### Zebrafish care, tail amputation, injection and bone staining

Wild-type^3^ (TAB-5) and Casper^4^ (*mpv^17a9^; mitf^aw2^*) zebrafish were obtained from the Zebrafish International Resource Center (Eugene, OR) and maintained at the University of Richmond animal facility following standard husbandry protocols^5^. All protocols were reviewed and approved by the Institutional Animal Care and Use Committee. All experiments were performed in animals 4 to 6 months of age. For amputations, fish were fully anesthetized in 0.2 mg/ml Tricaine (pH 7.0) until unresponsive to touch, placed on an inverted glass petri dish covered with Parafilm, and a sterile blade was used to cut 40 mm away from the distal tip of the caudal fin. For injection, fish were anesthetized and weighed to standardize the mass of C-dots injected per body weight (*μ*g/g). Then, fish were positioned on a wet sponge under the microscope and injected with C-dot intraperitoneally or intravascularly using a 36G Nanofill microsyringe attached to an electronically controlled micropump (UMP3 UltraMicroPump, WPI), as previously described^6, 7^. After manipulations, fish were allowed to recover for 30 minutes and returned to the animal facility where they received standard care for the duration of the experiment.

Live staining of mineralized structures was done using Alizarin Complexone (ALC; Sigma-Aldrich, A-3882). First, fish were sedated in Tricaine solution in fish facility water (0.1 mg/ml; pH 7.0). Then, fish were stained in a solution of ALC (10 mg/ml) and Tricaine (0.1 mg/ml; pH 7.0), for 30 min. To remove excess dye following exposure, fish were quickly rinsed in three sequential washes of fresh facility water and allowed to recover from sedation for 60 min before collecting the caudal fins for histology.

#### Histology

For histological sectioning, fish were anesthetized and the regenerated fin was amputated at a site proximal to the original cut. For non-amputated fish, the cut was done 40 mm away from the distal tip of the caudal fin. Immediately after caudal fin collection, the fish was placed in a recovery tank and the fin was splayed out by floating it on a large drop of 4% paraformaldehyde in 1x PBS inside a Petri dish. After a few minutes, once the fin submerged in the droplet, the Petri dish was closed and the fin was fixed for 4 hours. The fixative was then removed and replaced with a 10% sucrose solution. The Petri dish was sealed and stored at 4°C overnight. Samples were processed within 5 days. Fins were embedded in Clear Frozen Section Compound (VWR), frozen, and sectioned at 5 microns in a Leica CM1520 cryostat. Sections were collected on positively charged slides (Globe Scientific), allowed to dry for 30 minutes and covered with mounting media (30% glycerol in 1x PBS, 5 mg/ml n-Propyl gallate, 2.5 *μ*g/ml DAPI). Slides were kept in the dark at 4°C and sections were imaged within one week.

#### Imaging and analysis

For imaging, zebrafish were anesthetized and placed on a glass petri dish with the caudal tail splayed to separate the fin rays. Whole fin images were acquired under brightfield and epi-fluorescent light using a Zeiss Discovery V.20 dissecting microscope and a Zeiss Axiocam MRc camera. Filter sets for Fluorescein (green; 488/525 nm) and Texas Red (red; 596/615 nm) were used to detect C-dots and ALC, respectively. For consistency across experiments and biological replicates, all fluorescent images were taken using time exposures of 1 and 2.5 sec. Images were processed using Zeiss AxioVision SE64 v4.9.1 and fluorescence quantification was done using ImageJ v1.52. Fin sections were imaged using an Olympus Fluoview mounted on a fully automated Olympus IX83 inverted confocal microscope, using the company’s imaging software package. Appropriate wavelengths were used to detect C-dots and ALC. Representative images were cropped and assembled into figures using Adobe Photoshop v21.2.

Quantification of C-dots deposition on bones was based on the intrinsic photoluminescent properties of C-dots^1^. Fluorescent images of control and experimental fins were converted to 8-bit grayscale images using ImageJ, and the pixel intensities in areas of interest were obtained. In both control and experimental conditions, intensity values in regenerating areas were subtracted from background levels from non-regenerating regions. Then, intensity values in experimental conditions were normalized relative to control conditions. The integrated intensities for each region were averaged to determine relative fluorescence in arbitrary units for each treatment group. For signal profile analysis, the scale was set to 1 pixel = 1 *μ*m and a 1,300 *μ*m straight line was drawn starting 300 *μ*m anterior to the cut site along the 4th ventral-most ray. The signal profile was generated using the Analyze>Plot>Profile function of ImageJ. Signal intensity in regions of interest or along the profile line were recorded and analyzed in Excel. Graphical visualization of data was done in Excel or RStudio (code for signal intensity analysis in supplemental information).

## REFERENCES

1. Dan, Q.; Xiayan, W.; Yuping, B.; Zaicheng, S. Recent advance of carbon dots in biorelated applications. J. Phys. Mater. 2020, 3, 022003.

2. Zhou, Y.; Mintz, K. J.; Sharma, S. K.; Leblanc, R. M. Carbon Dots: Diverse Preparation, Application, and Perspective in Surface Chemistry. Langmuir 2019, 35, 9115–9132.

3. Mintz, K. J.; Zhou, Y.; Leblanc, R. M. Recent development of carbon quantum dots regarding their optical properties, photoluminescence mechanism, and core structure. Nanoscale 2019, 11, 4634–4652.

4. Yan, F.; Sun, Z.; Zhang, H.; Sun, X.; Jiang, Y.; Bai, Z. The fluorescence mechanism of carbon dots, and methods for tuning their emission color: a review. Mikrochim Acta 2019, 186, 583.

5. Zhou, Y.; Zahran, E. M.; Quiroga, B. A.; Perez, J.; Mintz, K. J.; Peng, Z.; Liyanage, P. Y.; Pandey, R. R.; Chusuei, C. C.; Leblanc, R. M. Size-Dependent Photocatalytic Activity of Carbon Dots with Surface-State Determined Photoluminescence. Appl Catal B 2019, 248, 157–166.

6. Zhou, Y.; Liyanage, P. Y.; Devadoss, D.; Rios Guevara, L. R.; Cheng, L.; Graham, R. M.; Chand, H. S.; Al-Youbi, A. O.; Bashammakh, A. S.; El-Shahawi, M. S.; Leblanc, R. M. Nontoxic amphiphilic carbon dots as promising drug nanocarriers across the blood-brain barrier and inhibitors of beta-amyloid. Nanoscale 2019, 11, 22387–22397.

7. Zheng, M.; Liu, S.; Li, J.; Qu, D.; Zhao, H.; Guan, X.; Hu, X.; Xie, Z.; Jing, X.; Sun, Z. Integrating oxaliplatin with highly luminescent carbon dots: an unprecedented theranostic agent for personalized medicine. Adv Mater 2014, 26, 3554–3560.

8. Sun, Y. P.; Zhou, B.; Lin, Y.; Wang, W.; Fernando, K. A.; Pathak, P.; Meziani, M. J.; Harruff, B. A.; Wang, X.; Wang, H.; Luo, P. G.; Yang, H.; Kose, M. E.; Chen, B.; Veca, L. M.; Xie, S. Y. Quantum-sized carbon dots for bright and colorful photoluminescence. J Am Chem Soc 2006, 128, 7756–7767.

9. Peng, Z.; Ji, C.; Zhou, Y.; Zhao, T.; Leblanc, R. M. Polyethylene glycol (PEG) derived carbon dots: Preparation and applications. Applied Materials Today 2020, 20, 100677.

10. Thakur, M.; Pandey, S.; Mewada, A.; Patil, V.; Khade, M.; Goshi, E.; Sharon, M. Antibiotic conjugated fluorescent carbon dots as a theranostic agent for controlled drug release, bioimaging, and enhanced antimicrobial activity. J Drug Deliv 2014, 2014, 282193.

11. Liyanage, P. Y.; Zhou, Y.; Al-Youbi, A. O.; Bashammakh, A. S.; El-Shahawi, M. S.; Vanni, S.; Graham, R. M.; Leblanc, R. M. Pediatric glioblastoma target-specific efficient delivery of gemcitabine across the blood-brain barrier via carbon nitride dots. Nanoscale 2020, 12, 7927–7938.

12. Iannazzo, D.; Ziccarelli, I.; Pistone, A. Graphene quantum dots: multifunctional nanoplatforms for anticancer therapy. J Mater Chem B 2017, 5, 6471–6489.

13. Li, X.; Vinothini, K.; Ramesh, T.; Rajan, M.; Ramu, A. Combined photodynamic-chemotherapy investigation of cancer cells using carbon quantum dot-based drug carrier system. Drug Deliv 2020, 27, 791–804.

14. Misra, S. K.; Ohoka, A.; Kolmodin, N. J.; Pan, D. Next generation carbon nanoparticles for efficient gene therapy. Mol Pharm 2015, 12, 375–385.

15. Li, S.; Amat, D.; Peng, Z.; Vanni, S.; Raskin, S.; De Angulo, G.; Othman, A. M.; Graham, R. M.; Leblanc, R. M. Transferrin conjugated nontoxic carbon dots for doxorubicin delivery to target pediatric brain tumor cells. Nanoscale 2016, 8, 16662–16669.

16. Wright, N. C.; Looker, A. C.; Saag, K. G.; Curtis, J. R.; Delzell, E. S.; Randall, S.; Dawson-Hughes, B. The recent prevalence of osteoporosis and low bone mass in the United States based on bone mineral density at the femoral neck or lumbar spine. J Bone Miner Res 2014, 29, 2520–2526.

17. Marie, P. J.; Kassem, M. Osteoblasts in osteoporosis: past, emerging, and future anabolic targets. Eur J Endocrinol 2011, 165, 1–10.

18. Gruneboom, A.; Kling, L.; Christiansen, S.; Mill, L.; Maier, A.; Engelke, K.; Quick, H. H.; Schett, G.; Gunzer, M. Next-generation imaging of the skeletal system and its blood supply. Nat Rev Rheumatol 2019, 15, 533–549.

19. Chen, G.; Qiu, H.; Prasad, P. N.; Chen, X. Upconversion nanoparticles: design, nanochemistry, and applications in theranostics. Chem Rev 2014, 114, 5161–5214.

20. Jung, J. S.; Jo, D.; Jo, G.; Hyun, H. Near-Infrared Contrast Agents for Bone-Targeted Imaging. Tissue Eng Regen Med 2019, 16, 443–450.

21. Li, S.; Skromne, I.; Peng, Z.; Dallman, J.; Al-Youbi, A. O.; Bashammakh, A. S.; El-Shahawi, M. S.; Leblanc, R. M. “Dark” carbon dots specifically “light-up” calcified zebrafish bones. J Mater Chem B 2016, 4, 7398–7405.

22. Peng, Z.; Miyanji, E. H.; Zhou, Y.; Pardo, J.; Hettiarachchi, S. D.; Li, S.; Blackwelder, P. L.; Skromne, I.; Leblanc, R. M. Carbon dots: promising biomaterials for bone-specific imaging and drug delivery. Nanoscale 2017, 9, 17533–17543.

23. Tonelli, F.; Bek, J. W.; Besio, R.; De Clercq, A.; Leoni, L.; Salmon, P.; Coucke, P. J.; Willaert, A.; Forlino, A. Zebrafish: A Resourceful Vertebrate Model to Investigate Skeletal Disorders. Front Endocrinol 2020, 11, 489.

24. Busse, B.; Galloway, J. L.; Gray, R. S.; Harris, M. P.; Kwon, R. Y. Zebrafish: An Emerging Model for Orthopedic Research. J Orthop Res 2020, 38, 925–936.

25. Witten, P. E.; Harris, M. P.; Huysseune, A.; Winkler, C. Small teleost fish provide new insights into human skeletal diseases. Methods Cell Biol 2017, 138, 321–346.

26. Salhotra, A.; Shah, H. N.; Levi, B.; Longaker, M. T. Mechanisms of bone development and repair. Nat Rev Mol Cell Biol 2020.

27. Rauch, F. Bone growth in length and width: the Yin and Yang of bone stability. J Musculoskelet Neuronal Interact 2005, 5, 194–201.

28. Suniaga, S.; Rolvien, T.; Vom Scheidt, A.; Fiedler, I. A. K.; Bale, H. A.; Huysseune, A.; Witten, P. E.; Amling, M.; Busse, B. Increased mechanical loading through controlled swimming exercise induces bone formation and mineralization in adult zebrafish. Sci Rep 2018, 8, 3646.

29. White, R. M.; Sessa, A.; Burke, C.; Bowman, T.; LeBlanc, J.; Ceol, C.; Bourque, C.; Dovey, M.; Goessling, W.; Burns, C. E.; Zon, L. I. Transparent adult zebrafish as a tool for in vivo transplantation analysis. Cell Stem Cell 2008, 2, 183–189.

30. Hoyte, D. A. N. Alizarin as an indicator of bone growth. J. Anat. 1960, 94, 432–442.

## References

1. Li, S.; Skromne, I.; Peng, Z.; Dallman, J.; Al-Youbi, A. O.; Bashammakh, A. S.; El-Shahawi, M. S.; Leblanc, R. M. “Dark” carbon dots specifically “light-up” calcified zebrafish bones. J Mater Chem B 2016, 4, 7398–7405.

2. Li, S. H.; Wang, L. Y.; Chusuei, C. C.; Suarez, V. M.; Blackwelder, P. L.; Micic, M.; Orbulescu, J.; Leblanc, R. M. Nontoxic Carbon Dots Potently Inhibit Human Insulin Fibrillation. Chem Mater 2015, 27, 1764–1771.

3. LaFave, M. C.; Varshney, G. K.; Vemulapalli, M.; Mullikin, J. C.; Burgess, S. M. A defined zebrafish line for high-throughput genetics and genomics: NHGRI-1. Genetics 2014, 198, 167–170.

4. White, R. M.; Sessa, A.; Burke, C.; Bowman, T.; LeBlanc, J.; Ceol, C.; Bourque, C.; Dovey, M.; Goessling, W.; Burns, C. E.; Zon, L. I. Transparent adult zebrafish as a tool for in vivo transplantation analysis. Cell Stem Cell 2008, 2, 183–189.

5. Westerfield, M., The zebrafish book: A guide for the laboratory use of zebrafish (Brachydanio rerio). 3rd edition ed.; University of Oregon Press: 1995.

6. Kinkel, M. D.; Eames, S. C.; Philipson, L. H.; Prince, V. E. Intraperitoneal injection into adult zebrafish. J Vis Exp 2010, 42, 2126.

7. Pugach, E. K.; Li, P.; White, R.; Zon, L. Retro-orbital injection in adult zebrafish. J Vis Exp 2009, 34, 1645.

